# Multicenter preclinical validation of next-generation CAR T cells: a strategy for harmonization, reproducibility, and its feasibility in clinical translation

**DOI:** 10.64898/2026.04.10.717659

**Authors:** Iman Dalloul, Markus Barden, Juliane C. Wilcke, Sophie Bernhard, Nicole Ellenbach, Anne-Laure Boulesteix, Hinrich Abken, Sebastian Kobold

**Author notes:** Contributed equally (Iman Dalloul, Markus Barden, Juliane C. Wilcke). Corresponding author: Sebastian Kobold, M.D., Institute of Clinical Pharmacology, Klinikum der Universität München, Lindwurmstraße 2a, 80337 Munich.

## Abstract

**Purpose:** Clinical translation of CAR T cell therapies has accelerated, yet preclinical evidence still often originates from single-center studies lacking sufficient robustness. Preclinical confirmatory multicenter studies have been proposed to improve the translational success, but their feasibility in cellular therapies remains unexplored.

**Methods:** We performed a confirmatory multicenter study validating C-C-motive-receptor-8 (CCR8) overexpression in CAR T cells—a strategy previously shown to enhance solid tumor infiltration. In vitro experiments covering activation, cytotoxicity, and migration using three CAR constructs were conducted across two centers with harmonized materials, preregistered protocols, randomization, and blinding.

**Results:** The data from the two centers confirmed key findings of the exploratory study: CCR8 overexpression in anti-EpCAM and anti-mesothelin CAR T cells leads to enhanced selective migration towards a CCL1-gradient, while not compromising antigen-specific T cell activatory capacity and cytotoxicity in vitro. The study furthermore broadened the applicability of CCR8 overexpression to anti-CEA CAR T cells.

**Conclusions:** This first-of-its-kind preclinical confirmatory CAR T study demonstrates the feasibility of a multicenter confirmation in cellular therapy, with technical and logistical challenges resolved through transparent communication between all parties involved. Both exploratory and confirmatory studies aim to downselect CAR candidates with the highest clinical success potential, as they compete for limited resources in preclinical research. It is therefore mandatory to clarify the extent of replications required to validate the experimental methodology and identify CAR candidates with most likelihood of success.

**TRANSLATIONAL RELEVANCE:** Preclinical evidence for novel CAR T cell therapeutic strategies relies mostly on exploratory single-center studies lacking robustness, with recent findings substantiating their limited predictive value for cellular therapies tested outside hematology. Here, the function of CCR8-armored CARs in vitro was confirmed in a preclinical confirmatory multicenter study, demonstrating the feasibility of such studies in adding value to the transition of preclinical concepts to clinical development. Our first-of-its-kind study may contribute to define new routes for preclinical testing and further raises the general question of what level of preclinical evidence is reasonably achievable in an academic context. It indicates the need for strong collaborative efforts to realize dedicated preclinical infrastructure for clinical translation of reprogrammed immune cellular therapeutics.

## INTRODUCTION

Within the last decade, chimeric antigen receptor (CAR) T cell therapy has revolutionized the treatment of certain hematological malignancies. The U.S. Food and Drug Administration (FDA) has approved seven CAR T cell products exclusively targeting hematological entities derived from the B cell lineage (1). Today, a wide array of CAR T cell products targeting other malignancies have now reached the stage of clinical evaluation. Currently, 2,447 interventional CAR T cell trials are registered at clinicaltrials.gov, of which 991 are actively recruiting (search term: CAR T cell; date:20-03-2026), underpinning the massive interest in these therapies. Despite such significant activities, extending the success of CAR T cell therapy to solid tumors has so far been anecdotal. Resistance of solid tumors to CAR T cell therapy can be caused by tumor heterogeneity (2, 3), the immunosuppressive tumor microenvironment (4, 5), and low infiltration into tumor tissue (6, 7).

We and others demonstrated that forced chemokine receptor expression is a promising strategy to enhance migration and infiltration of CAR T cells into the tumor tissue. In such a setting, general understanding of chemokine profiles in the tumor microenvironment is crucial in selecting appropriate chemokine axes to leverage for improving CAR T cell trafficking (6). We previously showed that arming CAR T cells with CXCR6 enhances the recruitment of genetically engineered T cells to cancer tissues expressing CXCL16 (8). We further probed the CCL1-CCR8 axis for tumor-directed recruitment, leveraging a chemokine axis otherwise employed by regulatory T cells and explored the concept in syngeneic and xenograft tumor models (9).

While these promising results illustrate the potential of the strategy for clinical application, we acknowledge the limitations of exploratory single-center studies. Here, the robustness of findings is mostly compromised by low sample sizes and high rates of false outcomes (10, 11). Along these lines, replicability of preclinical research in cancer biology is often frighteningly low (12–14), explaining in part the frequent failures in clinical translation. In the context of cellular therapies, this is of particular relevance. Recent evidence has demonstrated that while current single center models have their utility for proof-of-concept, they do not have predictive value for clinical activity beyond hematology and are thus unsuited for use in filtering the multitude of novel therapeutic concepts emerging from research (15). To increase chances of successful translation from bench to bedside, implementation of multicenter experimentation as performed in clinical trials has been suggested as a promising measure (16–19). With the considerable complexity of cellular therapeutics in general and of CAR T cells in particular, *confirmatory* preclinical multicenter trials, once successfully implemented, are thought to have great potential to substantiate knowledge claims from exploratory studies and guide decision making towards translation to clinical trials (10). Apart from these suggestions, the approach is globally still in its infancy and untapped in the context of cell therapy development.

Here we report the first multicenter preclinical confirmatory study in the field of cellular therapeutics. We aimed at confirming our previous results (9), investigating the value of CCR8 in improving the anti-tumor capacity of CAR T cells against solid tumors in both a murine (anti-EpCAM CAR) and a human model (anti-mesothelin CAR). The experiments were conducted in vitro with a focus on evaluating activation, cytotoxicity, and migration of T cells in a standardized manner. In addition, we aimed to extend our findings to another human model (anti-CEA CAR) in order to broaden the scope of applicability of CCR8 overexpression in CAR T cell therapy for solid tumors and to further increase the robustness of evidence for the success in clinical translation.

## MATERIALS AND METHODS

### Cell lines and virus production

Murine Panc02-EpCAM and human SUIT-mesothelin target cell lines have previously been engineered to express the respective target antigen (20); the LS174T (ATCC-CL188, RRID:CVCL_1384) cell line endogenously expresses CEA. 293Vec-Galv, 293Vec-Eco, and 293Vec-RD114 packaging cell lines were kindly provided by Manuel Caruso, CHU de Québec-Université Laval, Québec, Canada (21). To generate cell lines for stable production of virus containing the study-specific construct DNA, packaging cell lines were transduced with pMP71 retroviral vectors, kindly supplied by C. Baum, MHH Hannover, Germany. Murine 293Vec-Eco cells were engineered for GFP, anti-EpCAM CAR, and CCR8- anti-EpCAM CAR virus production, while human 293Vec-RD114 cells were engineered for GFP, anti-mesothelin CAR, CCR8-anti-mesothelin CAR, anti-CEA CAR, and CCR8-only virus production, as previously described (9). Target cell lines and producer cell lines were cultured in DMEM (Gibco, Cat. No. 41965-039) supplemented with 10 % (v/v) heat inactivated fetal bovine serum (FBS) (Gibco, Cat. No. 10270-106 / PAN-Biotech, Cat. No. P40-37500), 100 U/ml penicillin, 100 μg/ml streptomycin (Sigma Aldrich, Cat. No. P4333), and 2 mM L-glutamine (Sigma Aldrich, Cat. No. G7513).

### Retroviral transduction of murine and human T cells

For transduction of mouse T cells, splenocytes were isolated from C57Bl/6 mice (commercially provided by Janvier, RRID:MGI:2159769) and seeded at a concentration of 2 x 10^6^ cells/ml in RPMI 1640 (Gibco, Cat. No. 31870-025) supplemented with 10 % (v/v) heat inactivated FBS, 2 mM L-glutamine (Sigma Aldrich, Cat. No. G7513), 100 U/ml penicillin, 100 μg/ml streptomycin (Sigma Aldrich, Cat. No. P4333), 1 % (w/v) sodium pyruvate (Sigma Aldrich, Cat. No. S8636), 0.5 % (w/v) HEPES (Sigma Aldrich, Cat. No. H0887), and 50 μM β-mercaptoethanol (CarlRoth, Cat. No. Art.-Nr. 4227.3). Mouse T cells were activated for one day before retroviral transduction with anti-CD3e (Invitrogen, Cat. No. 16-0031-86, RRID: AB_2932551) and anti-CD28 monoclonal antibodies (Invitrogen, Cat. No. 16-0281-86, RRID:AB_467189) and 180 U/ml of IL-2 (Clinigen, Cat. No. M001692). On the day of transduction, retrovirus containing supernatant of the respective producer cell lines was added onto a 24-well plate coated with 12 µg/ml of retronectin (TAKARA, Cat. No. T100B) and centrifuged for two hours at 3,000 x g. Virus supernatant was removed and 10^6^ activated T cells stimulated with CD3/CD28 T cell activation Dynabeads (Gibco by ThermoFisher Scientific, Cat. No. 11453D) were added in each well. The next day, fresh medium supplement with 20 ng/ml IL-15 (Miltenyi Biotech, Cat. No. 130-095-766) was added to each well. For further expansion, T cell culture medium was replaced every second day and supplemented with IL-15. T cells were maintained at a concentration of 10^6^ cells/ ml.

For transduction of human T cells, peripheral blood mononuclear cells (PBMCs) were isolated from healthy human donors by density gradient centrifugation. For human donors, we purchased buffy coats from pseudonymized healthy blood donors. PBMCs were frozen, split in half, and sent to both sites for further utilization. After thawing, human T cells were seeded at a concentration of 10^6^ cells/ml in RPMI 1640 (Gibco, Cat. No. 31870-025) supplemented with 2.5 % (v/v) heat inactivated human serum (Sigma Aldrich, Cat. No. H4522) or FBS, 100 U/ml penicillin, 100 μg/ml streptomycin (Sigma Aldrich, Cat. No. P4333), 2 mM L-glutamine (Sigma Aldrich, Cat. No. G7513), 1 % (w/v) sodium pyruvate (Sigma Aldrich, Cat. No. S8636), and 1 % (v/v) “MEM Non-essential Amino Acid Solution” (Sigma Aldrich, Cat. No. M7145). Before retroviral transduction, T cells were activated for two days with CD3/CD28 T-cell activation Dynabeads (Thermo Fisher Scientific, Cat. No. 11132D), 180 U/ml of recombinant human IL-2 (Clinigen, Cat. No. M001692), and 2 ng/ml of human IL-15 (Miltenyi Biotech, Cat. No. 130-095-766). For human T cells, the same protocol as for the murine transduction was followed, with the exception that virus supernatant was centrifuged for 1.5 hours. T cells were expanded every second day under supplementation of 180 U/ml of recombinant human IL-2 and 2 ng/ml of human IL-15 (22).

Transduction efficiency was assessed via flow cytometry. Anti-EpCAM CAR expression on the T cell surface was detected by co-expressed GFP. The anti-mesothelin CAR, containing a c-myc tag, was detected using a c-myc antibody (Miltenyi, Cat. No. 130-116-485). Anti-CEA CAR was detected by goat F(ab’)2 anti-human IgG-PE antibody (SouthernBiotech, Cat. No. 2043-09, RRID:AB_2795669). Murine CCR8 was detected using an anti-murine CCR8-APC antibody (BioLegend, Cat. No. 150310, RRID:AB_2629601). Human CCR8 in the CCR8-anti-mesothelin CAR construct (bi-cistronic construct) was detected using an anti-human CCR8-PE antibody (BioLegend, Cat. No. 360604, RRID:AB_2562614). To generate T cells expressing anti-CEA CAR together with human CCR8, a double transduction was performed to co-express CCR8 and the CAR on the T cell surface (monocistronic construct for each). Human CCR8 was co-expressed with GFP for detection reasons.

In the activation and cytotoxicity assays, we adjusted the percentage of CAR+ cells to equal levels in the CAR and CCR8-CAR conditions within each center by diluting the cell suspension with the higher CAR+ percentage by non-transduced T cells of the same donor, which had undergone the same activation procedure as the transduced T cells. For the migration assays, no such adjustment was necessary because all relevant transduction rates are taken into account when calculating the odds ratios for each treatment condition (see below).

### Activation assay

Activation capacity of T cells was assessed by IFN-γ release. For anti-EpCAM CAR, T cells and tumor cells were pipetted at an effector to target (E:T) ratio of 10 T cells to 1 tumor cell. Accordingly, 30,000 tumor cells were cocultured with 300,000 T cells per well for 24 h. For anti-mesothelin CAR and anti-CEA CAR, T cells and tumor cells were pipetted at an E:T ratio of 5:1. Accordingly, 30,000 tumor cells were cocultured with 150,000 T cells per well for 24 h (CEA CAR) or 48 h (mesothelin CAR). Thereafter, plates were centrifuged and supernatants collected for ELISA (OptEIA™ Mouse IFN-γ ELISA Set, Cat. No. 5555138, and OptEIA™ Human IFN-γ ELISA Set, Cat. No. 555142, BD, respectively). Optical densities measured with the microplate reader were transformed to IFN-γ concentrations according to the corresponding technical data sheet with the following exceptions. For murine IFN-γ, we utilized log-log regression analysis without first subtracting the zero standard from optical densities (too many of them would have had to be excluded as negative values), and for human IFN-γ, we used linear instead of log-log regression analysis (because of the clearly linear relationship in the pre-run data from both centers and hence the better fit of the regression line).

### Cytotoxicity assay

Cytotoxicity of T cells was assessed by XTT cell viability assay. For anti-EpCAM CAR T cells, 10,000 tumor cells were cocultured with 100,000 T cells per well for 48 h. For anti-mesothelin and anti-CEA CAR T cells, coculture was performed under same conditions as the activation assay described above. Thereafter, 150 µl of the total 200 µl supernatant per well were removed. Adherent tumor cells remained in the wells and were incubated at 37 °C and 5 % (v/v) CO_2_ with reagent solution containing 50 µl XTT reagent (1 mg/ml) (PanReac AppliChem, Cat. No. A2240) in RPMI 1640 medium w/o supplements, 1 µl phenazine methosulfate (1.25 mM) (MERK, Cat. No. P9625), and 50 µl RPMI 1640 medium w/o supplements per well. The dye intensity of formazan is proportional to the number of metabolically active cells and was read in a spectrophotometer after 60 min and 120 min at 450 nm with 650 nm as reference. Dye intensity values measured at 120 min were analyzed as none of the wells had reached maximum measurable dye intensity at that time. Cytotoxicity of target cells was calculated for each treatment-condition well as follows: lysis [%] = 100 – [(OD_E+T_ – OD_E_) / (OD_T_ – OD_medium_) × 100], where OD = optical density, E = relevant effector cell (T cells), T = relevant target cell (tumor cells), and medium = tumor medium. All ODs were means of n = 3 or n = 5 wells, except OD_E+T_ with n = 1 for calculating percentage of lysis in this well.

### Migration assay

Migration of T cells towards murine CCL1 (2 ng/ml) and human CCL1 (10 ng/ml) (Biolegend, Cat. No. 584802 and 582706, respectively), was assessed using 3 µm transwell plates for murine T cells and 5 µm transwell plates for human T cells, respectively (Merck Millipore, Cat. No. MAMIC3S10 and MAMIC5S10, respectively). 500,000 T cells per well were pipetted in the upper chambers of the transwell plates. Migration was measured either towards migration media (RPMI 1640 medium supplemented with 1 % (w/v) BSA) without CCL1 (medium-only condition) or migration media with CCL1 located in the lower chambers of the transwell plates. After 3.5 hours of incubation at 37°C, the amount of cells migrated to the lower wells was quantified by flow cytometry. More specifically, we measured the percentage of CAR+ T cells for CAR and CCR8-CAR conditions and percentage of GFP+ T cells for the GFP condition in each lower well. To detect the CCR8-CAR construct in anti-CEA CAR T cells, we first gated on CAR+ T cells and then on CCR8+ in this CAR+ population. Enrichment of migrated cells was calculated as the following odds ratio for each treatment-condition well: Odds ratio = Odds of treatment condition / Odds of corresponding medium-only condition, where Odds = % of migrated cells / (100 % – % of migrated cells). For each medium-only condition, we averaged the percentage of migrated cells across wells before determining the odds.

### Statistical analyses

All analyses were performed using R Statistical Software version 4.2.2 (23, RRID:SCR_001905), using one-sided testing of directional hypotheses with an alpha level of 0.05. We nevertheless use the convention of two-sided 95 % confidence intervals (CIs) around effect estimates.

Two-way between-group ANOVAs with the interaction effect included were used to analyze the data of each experiment (with type III sums of squares if unbalanced). The independent variables were treatment condition (CCR8-CAR, CAR, GFP) and research center (A, B). The dependent variable was IFN-γ release (in pg/ml) in the activation experiments, tumor cell lysis (in %) in the cytotoxicity experiments, and the enrichment of the migrated cells expressed as an odds ratio in the migration experiments. Planned contrasts were used for all comparisons between two groups, after extreme outliers had been excluded (defined as 3 times the interquartile range above the third quartile or below the first quartile within a cell of the 3 x 2 design; overall 3 % of the data were excluded; sensitivity analyses with the complete data sets were additionally performed to check the robustness of the results) and after the planned analyses had been inferred to be robust against observed violations of the test assumptions (we simulated obvious failures of normality in a design cell, apparent in QQ plots, histograms or Shapiro-Wilk tests, or heterogeneity of variance between design cells in unbalanced data, apparent in residual plots or the Levene test, based on our data to check the robustness of the ANOVAs in terms of type 1 error and power).

In the cytotoxicity experiments, tests of non-inferiority were performed with a non-inferiority margin of 10 % of the arithmetic mean of the CAR condition, and the *p* values of the two hypothesis tests were adjusted for multiple comparisons using the Holm correction. We deemed this correction necessary because either significant result on its own would already be interpreted as showing the efficacy of the CCR8-CAR condition (24). No such correction was used in the other experiments, either because there was only one confirmatory hypothesis or because the hypotheses were not considered equivalent (the secondary hypothesis does not address the main comparison of interest here, i.e., the migratory superiority of CCR8-CAR over CAR T cells). For each confirmatory hypothesis, we checked that the interpretation of the main effect of interest need not be qualified because of a significant disordinal or hybrid treatment-by-center interaction. CIs were calculated for the marginal means corresponding to the confirmatory hypothesis tests in the ANOVA models, using the corresponding standard errors. For easier interpretation of the effect estimates in the migration experiments, we give the odds ratios of interest (instead of their differences to the empirical odds ratios of the two negative control conditions, as in the hypothesis tests).

The number of wells in each experiment was determined to obtain at least 0.90 power for detecting the effects specified in the hypotheses (and preferably also sufficient power for detecting any treatment-center interactions relevant for their interpretation) as well as sufficiently reliable results and reasonably accurate effect size estimates for the tested settings, using the R package Superpower (25), based on pilot data generated during the planned pre-runs for harmonizing procedures between centers. Each treatment condition had 10 wells per experiment and center, except for 17 wells in the EpCAM cytotoxicity assay. All other conditions had 5 wells per center, and we used 3 ELISA standards per center; for EpCAM, these numbers were 3 and 2, respectively (cf. plate layouts). More details on the sample size calculation can be found in the preregistrations and analysis codes on the OSF platform (access upon reasonable request to the corresponding author).

## RESULTS

### Design of the preclinical confirmatory multicenter study assessing CCR8-CAR T cell function

The purpose of this confirmatory multicenter study was to test the overall hypothesis that CCR8 improves the anti-cancer capacity of CAR T cells across two research centers with an independent data analysis team. Figure 1 gives an overview of the experiments and overall procedures in the study. We employed in vitro model systems to investigate activation, killing, and migration capacities of CCR8-CAR T cells targeting the solid-cancer antigens EpCAM (murine), mesothelin (human), and carcino-embryonic antigen (CEA) (human). The study was jointly designed by the two centers performing the experiments, the Division of Clinical Pharmacology at LMU Hospital Munich, Germany, and the Division of Genetic Immunotherapy at the Leibniz Institute for Immunotherapy (LIT) in Regensburg, Germany, in cooperation with the Institute for Medical Information Processing, Biometry, and Epidemiology (IBE) at LMU Munich, Germany, which was responsible for the statistical analysis plan, preregistrations, and data analysis.

**Figure 1:**
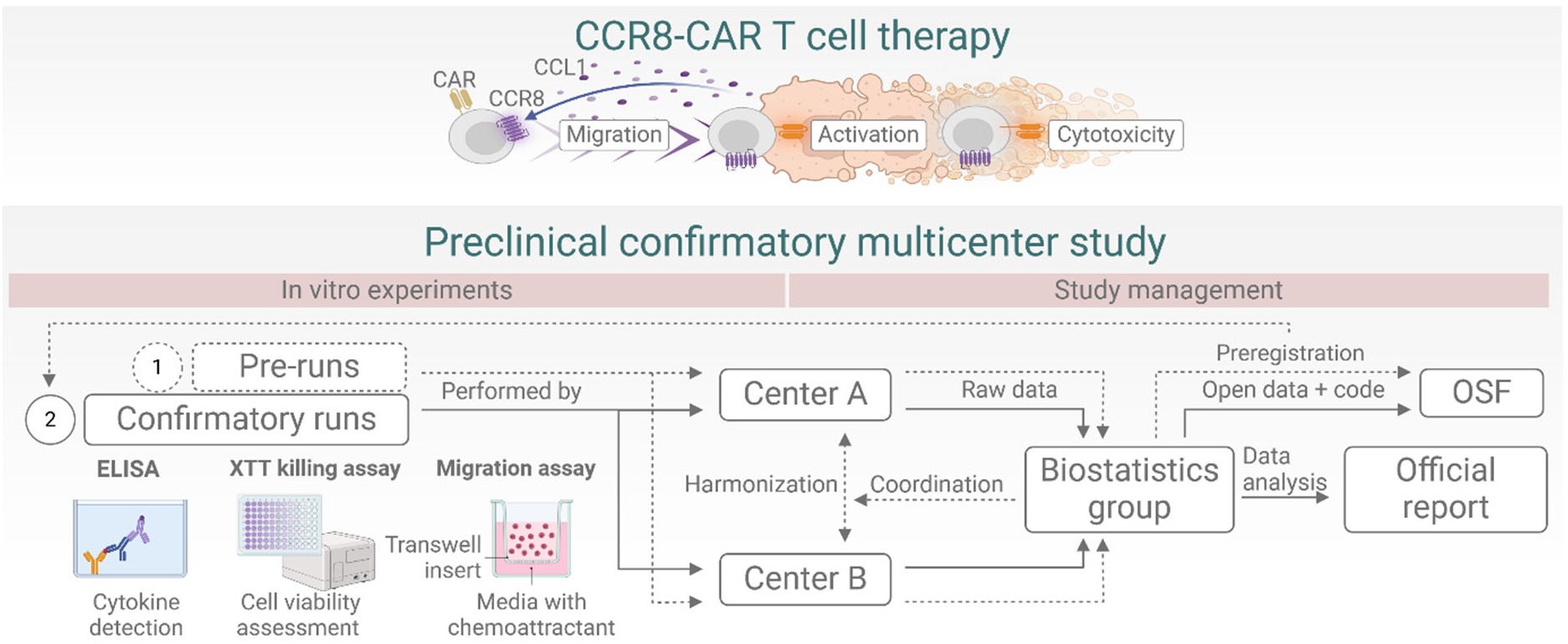
Study organigram. Activation, cytotoxicity, and migration of CCR8-CAR T cells were assessed in a preclinical confirmatory multicenter study in the centers A and B. Pre-runs were performed to harmonize the procedures between the centers in coordination with the biostatistics group. Consecutively, confirmatory runs were preregistered on the Open Science Framework (OSF) platform and performed in both centers by blinded experimenters. Raw data were analyzed by the biostatistics group and prepared for official report, with open data and code made available via OSF. Graphics was created in BioRender. Abken, H. (2026) https://BioRender.com/u4kj6jj.

All experiments in our study had a fully crossed 3 x 2 between-group design with the factor treatment (one experimental and two control conditions) and the factor research center (two fixed levels). CCR8 and CAR transduced T cells (CCR8-CAR) were the experimental condition of interest. CAR transduced T cells (CAR) were used as a control for CAR-related background activity in the activation and cytotoxicity experiments or to control for migration of CAR T cells. GFP-transduced T cells were used as a control for general T cell background activity, cytotoxicity, or migration. In the activation and cytotoxicity experiments, GFP-transduced T cells were used as negative control, whereas CAR-only transduced T cells were used as positive control. A tumor-only condition was added as qualitative control. In migration experiments, the former two conditions were both used as negative control for CCR8 function.

Prior to confirmatory experiments, preliminary trials (pre-runs) were conducted in both centers to harmonize and standardize protocols and to determine required sample sizes. To optimize experimental parameters, pre-runs were initially conducted in a single center by testing different E:T ratios, different incubation durations (24 h, 48 h, and 72 h) for ELISA and XTT assays, and different concentrations of CCL1 (2 ng/ml and 10 ng/ml) for migration assays. For each experiment, one or a few best parameter settings were identified and implemented in the corresponding pre-run in the other center. Final parameter settings and sample sizes were then determined based on the pre-run data from both centers.

After analyzing pre-run data and before performing confirmatory runs, each experiment was preregistered with the selected conditions on the Open Science Framework (OSF) platform, along with the confirmatory hypotheses, study design, and planned analyses, in order to increase transparency, credibility, and reproducibility of the research. Based on our experience with the pre-runs and other studies, we also specified a number of conditions for excluding data from an experiment in one center and repeating the experiment in that center. These concerned aspects such as the minimum transduction efficiency, number of available CAR T cells and their viability, implausible optical densities of T cell-only conditions compared to the tumor-only condition (cytotoxicity assay), irregular growth of tumor cells in the tumor-only condition (cytotoxicity and activation assays), and manipulation check failures. The preregistrations of all experiments as well as the data sets of all confirmatory findings, plate layouts, analysis codes, and deviations from the preregistrations are accessible upon reasonable request to the corresponding author.

Experimenters and data analysts were blinded to the treatment conditions during experimental procedures and data analysis of all confirmatory experiments to eliminate any potential bias and enhance the internal validity, robustness, and objectivity of the research outcome. Blinding was achieved by grouping the wells in the 96-well-format microtiter plates according to a predetermined layout, assigning these groups to three treatment conditions in one center using a simple randomization procedure in the statistical software R, and counterbalancing their order in the other center (only the order of the mesothelin migration assay was not counterbalanced, in order to maintain blinding in the center that had to repeat the experiment after a staining failure). Unblinding was only performed after confirmatory hypothesis tests had been completed.

### Multicenter data confirm that antigen-specific CAR T cell IFN-γ release is not attenuated by co-expression of CCR8

As a prerequisite for our overall hypothesis that CCR8 improves the anti-cancer capacities of CAR T cells against solid tumors, we investigated whether co-expression of CCR8 with the CAR alters basic CAR T cell activatory functions. CAR-mediated T cell activation capacity was assessed by IFN-γ ELISA in three CAR systems: murine anti-EpCAM CAR, human anti-mesothelin CAR, and human anti-CEA CAR. In each system, T cells equipped with the CAR only as well as T cells expressing both CAR and CCR8 were compared to GFP-transduced T cells, which functioned as negative control in the manipulation checks (CAR > GFP in each center) and the confirmatory hypothesis tests.

For activation experiments, we hypothesized based on previous data (9), that IFN-γ amounts released by CCR8 CAR T cells would be higher than those released by GFP T cells. Confirmatory-run data confirmed the superiority of CCR8-CAR T cells over GFP T cells across both centers in all three CAR systems (Fig. 2A-C; CCR8-CAR > GFP: *t*(54) = 32.84 for EpCAM, *t*(53) = 5.57 for mesothelin, *t*(52) = 12.93 for CEA, all p < 0.001). IFN-γ release of CCR8-CAR was higher than that of GFP by 477 pg/ml, 95 % CI [448, 506], for EpCAM, 59 pg/ml, [38, 80], for mesothelin, and 61 pg/ml, [51, 70], for CEA. Notably, variations in IFN-γ release were observed between the two centers, most prominently for CCR8-anti-EpCAM CAR (Fig. 2A). The potential center-specific influences on CAR T cell behavior were, however, qualitatively different for each activation experiment.

**Figure 2.**
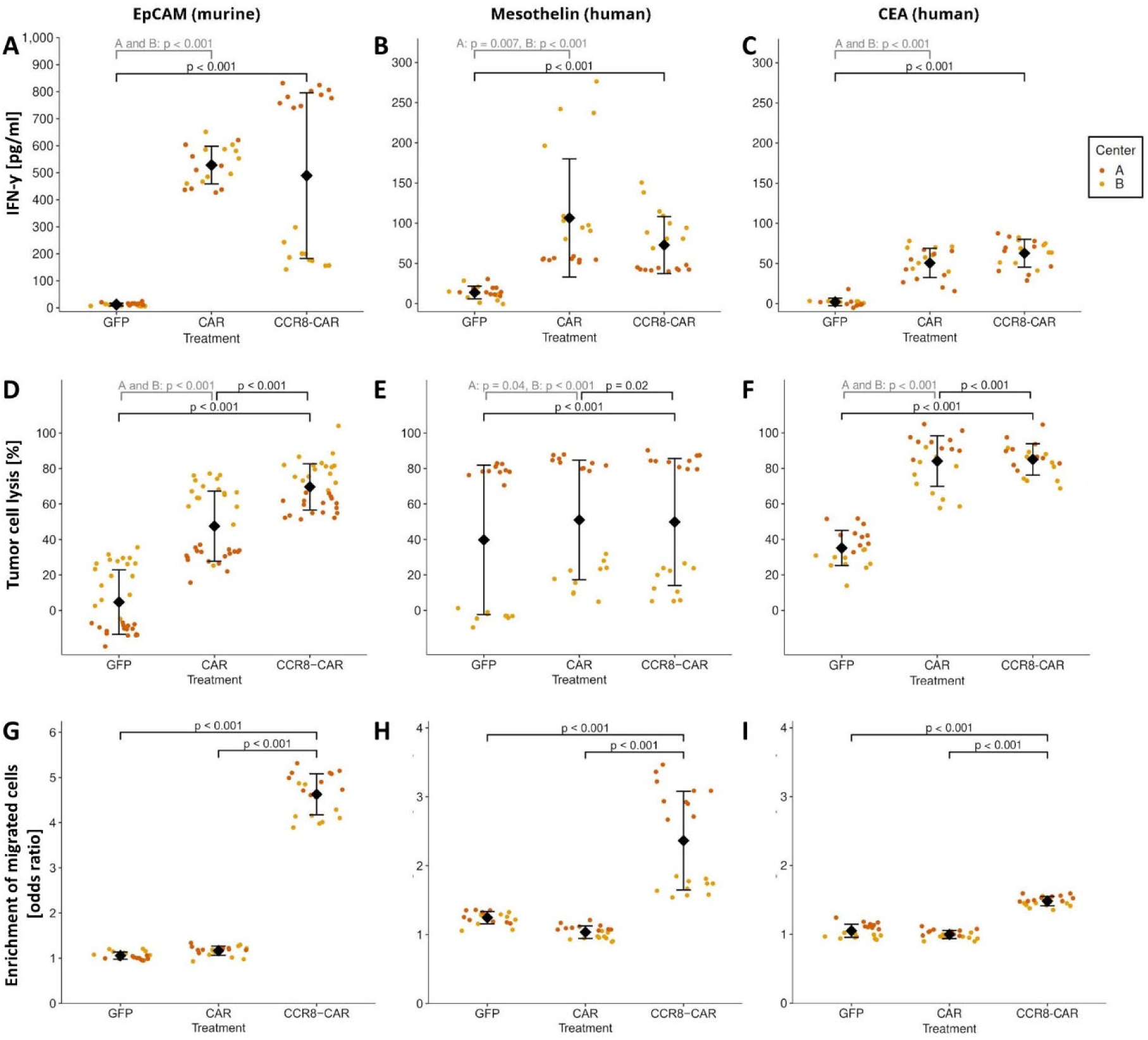
Multicenter validation of CAR T cell function: IFN-γ secretion, cytotoxicity, and migration. CAR T cell effector functions were assessed by ELISA recording IFN-ƴ, XTT-based viability assay, and transwell migration assay across two independent centers. (A–C) IFN-γ secretion measured after T cell coculture with Panc02-EpCAM cells (A, 24 h, 10:1 E:T ratio), SUIT-mesothelin cells (B, 48 h, 5:1 E:T ratio), or LS174T cells (C, 24 h, 5:1 E:T ratio); supernatants were diluted to 1:500 (EpCAM and CEA) or 1:200/1:1,000 (mesothelin, centers A/B). (D–F) Cytotoxicity was assessed after 48 h coculture with Panc02-EpCAM (D, 10:1 E:T), 48 h with SUIT-mesothelin (E, 5:1 E:T), or 24 h with LS174T (F, 5:1 E:T). (G–I) In vitro migration was recorded in a transwell system toward 2 ng/ml murine CCL1 (G, EpCAM) or 10 ng/ml human CCL1 (H, mesothelin; I, anti-CEA). Gray bars depict manipulation checks (A–F, CAR > GFP; G–I, odds ratio = 1 ± 0.25 for GFP and CAR), and black bars confirmatory test results (A–C, CCR8-CAR > GFP; D–F, CCR8-CAR > GFP and CCR8-CAR > CAR; G–I, primary: CCR8-CAR > CAR, secondary: CCR8-CAR > GFP). Mean ± standard deviation was calculated across both centers. Y-axis scaling differs between murine (A, G) and human models (B–C, H–I). Statistical details reported in main text.

### Multicenter data confirm that CAR T cell cytotoxicity is not attenuated by co-expression of CCR8

We further investigated whether co-expression of CCR8 with the CAR alters CAR T cell cytotoxic function. CAR-mediated cytotoxicity was assessed by XTT cell viability assay in the three CAR systems described above. In each system, T cells equipped with CAR only and CCR8-CAR T cells were compared to each other as well as to GFP-transduced T cells, which functioned as negative control in the manipulation checks (CAR > GFP in each center) and the confirmatory hypothesis tests.

For cytotoxicity experiments, we hypothesized that tumor cell lysis (%) mediated by CCR8-CAR T cells would be higher than lysis by GFP T cells, but not lower than lysis by CAR only T cells. In all three CAR systems, confirmatory-run data confirmed superiority of CCR8-CAR T cells to GFP T cells and non-inferiority of CCR8-CAR T cells to T cells expressing only the CAR across both centers (Fig. 2D-F; CCR8-CAR > GFP and CCR8-CAR ≥ CAR: *t*(94) = 38.13 and *t*(94) = 14.73 for EpCAM, *t*(53) = 6.68 and *t*(53) = 2.19 with p = 0.016 for mesothelin, *t*(52) = 19.58 and *t*(52) = 3.95 for CEA, all other p < 0.001; *p* values Holm-adjusted within each experiment). The difference between CCR8-CAR and GFP lysis was 64.2 %, 95 % CI [60.9, 67.6], for EpCAM, 12.3 %, [8.6, 15.9], for mesothelin, and 49.9 %, [44.8, 54.9], for CEA, and that between CCR8-CAR and CAR 20.2 %, [16.8, 23.5], for EpCAM, -1.1 %, [-4.8, 2.5], for mesothelin, and 0.9 %, [-4.1, 6.0], for CEA. Variations between centers were most pronounced in the anti-mesothelin CAR system with increased background cytotoxicity by GFP T cells in one center (Fig. 2E). Despite the absolute increase in cytotoxicity, we still detected a relative increase in cytotoxicity induced by the CAR.

### Multicenter data confirm that CCR8 increases in vitro migratory capacity of CAR T cells

Our overall hypothesis that CCR8 improves the anti-cancer capacities of CAR T cells against solid tumors relies on the functionality of the CCR8-CCL1 migratory axis. To evaluate the impact of CCR8 on migratory capacities of CAR T cells in vitro, we performed migration assays towards CCL1. In each system, T cells equipped with CCR8-CAR T cells were compared to CAR-only and to GFP-transduced T cells, which served as a form of manipulation check in each center (expected odds ratio ≍ 1) and as negative controls in the confirmatory hypothesis tests.

For migration experiments, we hypothesized that migration for CCR8-CAR T cells would be higher than for T cells expressing CAR only. Our secondary hypothesis was that migration for CAR-CCR8 is also higher than for GFP T cells. In all three CAR systems, confirmatory-run data confirmed that migration of CCR8-CAR T cells is higher compared to both CAR and GFP T cells across both centers (Fig. 2G-I; CCR8-CAR > CAR and CCR8-CAR > GFP: *t*(54) = 59.60 and *t*(54) = 61.48 for EpCAM, *t*(54) = 32.48 and *t*(54) = 27.33 for mesothelin, *t*(54) = 33.45 and *t*(54) = 29.79 for CEA, all p < 0.001). The CAR-CCR8 odds ratio was 4.63, 95 % CI [4.54, 4.71], for EpCAM, 2.36, [2.31, 2.42], for mesothelin, and 1.48, [1.46, 1.51], for CEA, and thus markedly higher than one (where migration is identical with and without the CCL1 ligand). Between-center-variation was lower in the migration experiments than in the activation and cytotoxicity experiments, affecting mainly the CCR8-anti-EpCAM and anti-mesothelin CARs (Fig. 2G-I), with stronger effects for both CCR8-CARs in the same center.

## DISCUSSION

We report results of a first-of-its-kind preclinical confirmatory multicenter study using CAR T cells (Fig. 3). Mechanistic advantages of CCR8 overexpression were confirmed in the context of CAR T cell functions in vitro in EpCAM- and mesothelin-expressing cancer cell models and were extended to a CEA-expressing cancer cell model. CCR8-CAR T cells retain adequate antigen-specific activatory strength. While their cytotoxic functions are similar to CAR-only T cells, CCR8-CAR T cells show a marked increase in migratory capacities. Notably, the observed variability between centers and inconsistency between assays did not affect our findings regarding the directional confirmatory hypotheses.

**Figure 3:**
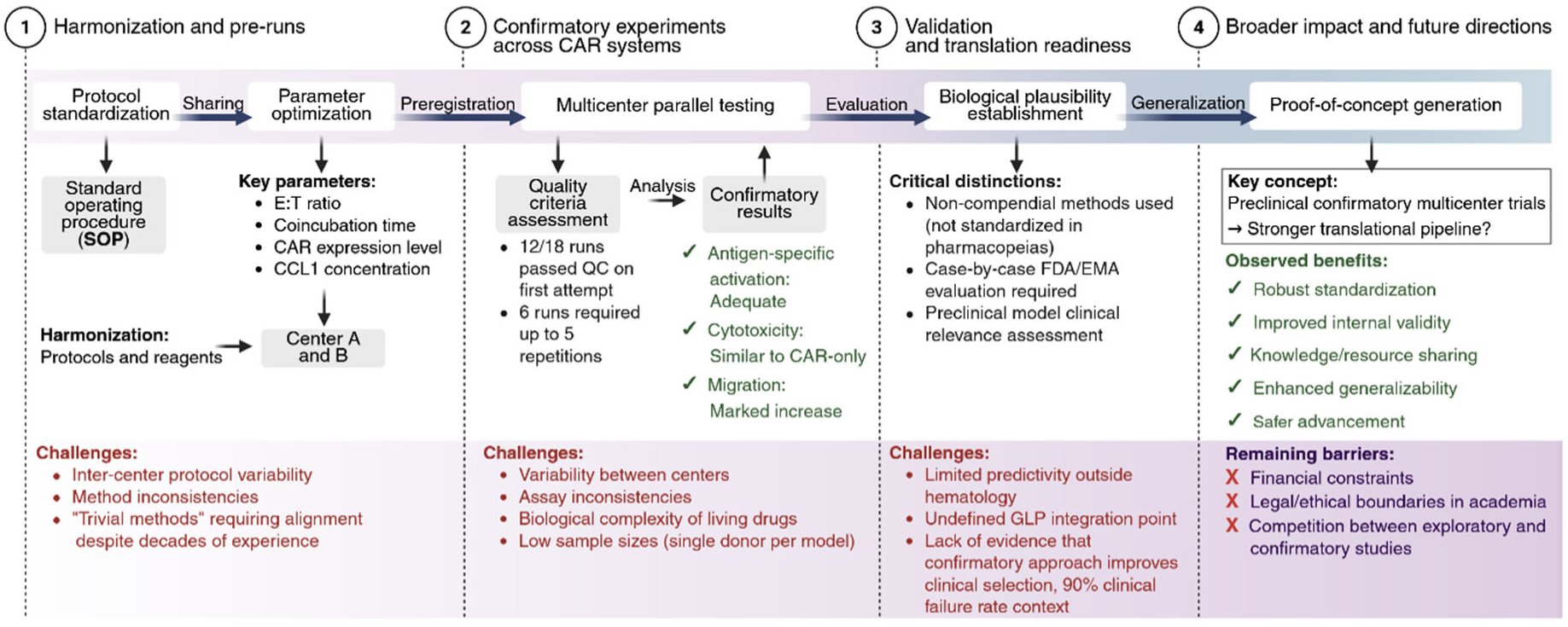
Key steps and learnings for translational cancer immunotherapy. Schematic illustrating how the preclincial confirmatory multicenter study approach was embedded into the core translational pathway from preclinical harmonization to clinical translation readiness in CAR T cell therapy for cancer, highlighting challenges, barriers, and observed benefits. Created in BioRender. Abken, H. (2026) https://BioRender.com/e4zpayd.

Pre-runs were essential for harmonizing and standardizing protocols (26) and contributed relevantly to the technical success of the confirmatory multicenter research study. Kinetics of T cell activation, cytotoxicity, and migration may vary dependent on unique characteristics of the respective CAR system, such as CAR and/or CCR8 expression levels on the T cell surface. Therefore, we established the main experimental parameters, such as effector T cell numbers or effector to target (E:T) ratio and coincubation time, for each of our three CAR systems independently by performing pre-runs prior to the confirmatory experiments. In addition, T cells and some reagents were shared between centers for the purpose of harmonization and economical experimentation. With pre-runs and standardization, 12 out of the overall 18 confirmatory experimental runs across both centers met the pre-defined quality criteria on the first attempt (cf. exclusion conditions in “Study Design”). However, the other six confirmatory experimental runs had to be repeated up to five times until they met the pre-defined quality criteria. This emphasizes once more the challenging nature of performing even a highly standardized confirmatory study in the biologically complex CAR T cell setting (27) and highlights the importance of pre-defined quality criteria (28, 29). Preclinical research can greatly benefit from such increased transparency regarding procedural challenges, which, although they regularly occur, are not commonly reported in exploratory research.

Limitations of the reported study further include its restriction to comparisons within CAR systems. Stepping up from an exploratory to a confirmatory setting was accompanied by increase in sample sizes, which was challenging to realize in terms of limited resources and automation. Given the probable overpowering of the in vitro experiments for the confirmatory hypotheses (cf. our rationale in the “Statistical Analysis”), it is important to consider whether the identified effects are large enough to be investigated further (30). Note, however, that the reported effect sizes are donor-dependent and, given that we used only one T cell donor per model for all pre-runs and confirmatory experiments in both centers, these cannot be generalized to other settings. With single wells as the experimental unit, the low variability between them increased the likelihood of achieving statistical significance. In an ideal scenario, more donors and cell preparations are preferred and the wells would only be treated as subsamples (31). Since this is a first-of-its-kind CAR T cell preclinical confirmatory multicenter study, we already solved many of the technical and logistical challenges through intensive transparent communication between all parties involved. We recently discussed our results and the limitations of preclinical confirmatory multicenter studies within the DECIDE meta-research project, which comprises seventeen such biomedical studies conducted in Germany. Analysis of the EPICYCLE study (32) and other unpublished studies involved, revealed recurrent methodological challenges in preclinical research. These common pitfalls were systematically evaluated and used to develop targeted strategies to improve study design, coordination, and conduct (33).

CAR T cells are “living drugs“, which present a new set of challenges for clinical translation (34). The U.S. Food & Drug Administration (FDA) and the European Medicines Agency (EMA) acknowledge the exceptional status of CAR T cells in drug development by categorizing CAR T cells as advanced therapy medicinal products (ATMPs) (35). CAR T cells may be seen as the most personalized embodiment of a therapy and thus prone to a high number of variables (36). More than 50 % of CAR T cell trials in the USA are sponsored by academia (37), highlighting the pivotal role of academia inthis field of clinical translation. The FDA recently published a guidance document for the development of CAR T cell products intended to assist industry and academic sponsors (38). Due to the biological complexity of CAR T cell products, the FDA recommends a case-by-case preclinical testing strategy to evaluate feasibility, safety, and efficacy of novel CAR T cell therapeutic modalities. Testing strategies for CAR T cell therapies include non-compendial methods (32), which are defined as methods not standardized in the US or EU Pharmacopeias.

The evidence level of preclinical data required as a basis for clinical translation of CAR T cell therapeutic concepts remains to be defined by a case-by-case evaluation through the respective authority. The primary objective of any preclinical assessment is the establishment of biological plausibility. According to guidelines by the EMA, proof of concept studies should generate non-clinical evidence supporting the potential clinical effect (39). In a first step of establishing a CAR T cell proof-of-concept study based on non-compendial methods, faithful preclinical models to evaluate novel CAR T cell therapies need to be identified (40). In the wide range of in vitro models available for testing CAR T cell safety and efficacy, it is key to select reliable models of high clinical relevance that may predict later clinical efficacy (41, 42). Current evidence from reported clinical testing in relation to preclinical testing shows the lack of predictivity of preclinical results beyond hematology for later clinical testing (15). While this is not to be mixed with the clear utility for proof-of-concept, additional models and testing routes need to be defined to allow refinement of concepts towards clinical testing. Multicenter validation may be one possible building block in this situation. In a consecutive step, clinically relevant models would need to be applied under quality-controlled conditions. It remains undefined at which point good laboratory practices (GLPs) should be integrated into the translational pipeline for cellular therapeutics. The GLP test facility of the Fraunhofer Institute for Cell Therapy and Immunology, Leipzig, Germany, is a prime example of how preclinical GLP-compliant studies may be realized in the academic context (43).

As of today, exploratory studies represent the main study format in preclinical CAR T cell assessment. The robustness of results from exploratory studies is, however, often attenuated by low sample sizes and false positive outcomes. The issue of preclinical findings not being easily reproducible together with drug developments failing to show clinical effectiveness is gaining attention in the field. As part of the “Reproducibility Project: Cancer Biology”, the variabilities in replicating preclinical cancer research from other research centers have been highlighted, underscoring the challenges of translating preclinical research into clinical applications (12). Similarly, Bayer and Amgen, two major industrial laboratories, state that they could only replicate 11 % and 25 % of the results, respectively, from several major cancer studies (13, 14). This lack of reproducibility is a major concern with respect to the robustness of early-stage research and finally the ability to turn those findings into effective treatments. When pre-clinical results cannot be reliably reproduced, it drives up the costs of drug development due to increased frequencies of failed attempts. However, an unsolved aspect is the weight of methodological challenges in replication across laboratories and centers (14, 44). Under such considerations, it will be critical to discern issues with the method employed versus the actual biological effect or mechanism, the latter of which is the scope of our study.

Preclinical confirmatory multicenter trials hold the promise to strengthen translation from basic to clinical research (10). The FDA gives advice for multisite testing, but so far only in the context of multisite manufacturing, recommending the use of the same SOPs, reference materials, reagents, and equipment across testing facilities (38). Adhering to similar principles, we herein applied a multicenter study approach at the initial step of preclinical in vitro CAR T cell testing, in order to confirm the potential of our CAR T cell therapy candidates before entering more laborious and costly stages of clinical translation. As described, we faced challenges in harmonizing protocols and seemingly trivial methods in two leading laboratories with decades-long experience in CAR T cell engineering and testing (45, 46). Only after successfully completing said harmonization, which critically was applied to the exact settings of the project, could we truly focus on the underlying biological effect. It is clear that more complex systems, such as advanced disease in vitro and in vivo models, require even more laborious harmonization efforts than applied here. Such effort, however, is impaired by financial, legal, and ethical boundaries, which are challenging in an academic setting and create a competition between exploratory and confirmatory studies. It is further questioned by the current absence of evidence that the confirmatory approach rolled out to all aspects of preclinical testing would really be able to improve selection and confidence in clinical testing. However, there is more consensus on the benefits of the multicenter aspect of preclinical studies such as robust standardization, improved internal validity, sharing knowledge and resources, enhanced generalizability of findings across laboratories, and faster advancement in translational research (16, 47–49). Bearing in mind the stunningly high 90 % failure rate of clinical drug development and the ever-tightening financial resources of public funding, the need to reduce costs and increase speed has never been higher (50). With this study, we leverage multidisciplinary networks specialized on preclinical confirmatory multicenter research, such as the DECIDE consortium, and commend concerted efforts to work towards enhancing reproducibility and clinical translation in oncology and beyond.

## DECLARATIONS

### Acknowledgements

This study was funded by the Federal Ministry of Education and Research (BMBF), grant 01KC2005 (CONTRACT to SK and HA), as well as the Bavarian Cancer Research Center (BZKF) (TANGO to SK), the Deutsche Forschungsgemeinschaft (DFG, grant numbers KO5055/2-1 and KO5055/3-1 to SK and grant BO3139/7-2 to ALB), the international doctoral program ‘i-Target: immunotargeting of cancer’ (funded by the Elite Network of Bavaria; to SK), the Melanoma Research Alliance (grant number 409510), Marie Sklodowska-Curie Training Network for Optimizing Adoptive T Cell Therapy of Cancer (funded by the Horizon 2020 programme of the European Union; grant 955575 to SK), Marie Sklodowska-Curie Training Network for tracking and controlling therapeutic immune cells in cancer (funded by the Horizon Programme of The EU, grant 101168810 to S.K.), Else Kröner-Fresenius-Stiftung (IOLIN to SK), German Cancer Aid (Deutsche Krebshilfe, AvantCAR.de, and 70117182 to SK), the Wilhelm-Sander-Stiftung (to SK), Ernst Jung Stiftung (to SK), Institutional Strategy LMUexcellent of LMU Munich (within the framework of the German Excellence Initiative; to SK), the Go-Bio-Initiative (to SK), the m4-Award of the Bavarian Ministry for Economical Affairs (to SK), Bundesministerium für Bildung und Forschung (Binostics to SK), European Research Council (Starting Grant 756017, CoG 101124203 and PoC Grant 101100460 to SK), by the SFB-TRR 338/1 2021–452881907 (to SK), Fritz-Bender Foundation (to SK), Deutsche José Carreras Leukämie Stiftung (to SK), Hector Foundation (to SK), Monika-Kutzner Foundation for Cancer Research (to SK), Bavarian Research Foundation (BAYCELLATOR to SK), the Bruno and Helene Jöster Foundation (360° CAR to SK) the Dr. Rurainski Foundation (to S.K.), and the Constanze and Dr. Brigitte Wegener Foundation (to S.K.).

We thank Anja Pavlica and Charlotte Schenkel for technical assistance, Kathrin Manske and Thaddäus Strzalkowski for support with the harmonization, and Luzia Hanßum for her assistance with preparing the R scripts. We also thank Pedro Mesquita. Flow cytometry data were collected at the Flow Cytometry Core Facility, University Hospital, LMU Munich, with a Beckman Coulter CytoFLEX. We also acknowledge the fruitful exchanges within and outside the DECIDE workshops organized by the QUEST Center for Responsible Research, Berlin Institute of Health (BIH) at Charité, Berlin, Germany, on different aspects of preclinical confirmatory multicenter studies.

### Author Contributions

ID, MB and SB performed or assisted with the experiments. JW and NE supported the project methodologically, including preregistrations, coordination, and data analyses. SK, HA and ALB supervised the project and provided the funding. ID, MB, JW and SK designed the experiments in detail and wrote the paper. All authors read and approved the final manuscript.

### Ethics approval

Blood from healthy donors was obtained under Ethics approvals 21-2224-101 (University Hospital Regensburg) and 22-0259 (LMU Munich).

